# Towards inferring atomic scale conformation landscape of biomolecules from cryo-electron tomography data

**DOI:** 10.64898/2026.02.15.706028

**Authors:** Florène Shahrzad Feyzi, Slavica Jonic

## Abstract

Understanding continuous conformational variability of biomolecular complexes at atomic resolution is essential for linking structure to function, but remains challenging for cryo-electron tomography (cryo-ET) data due to high noise and missing-wedge (MW) artifacts. Physics-based methods, such as MDTOMO (based on classical molecular dynamics simulations of a given atomic structure to flexibly fit subtomograms), provide accurate estimation of atomic coordinates from subtomograms, but their computational cost limits large-scale applications. We present DeepMDTOMO, a supervised deep learning (DL) framework that uses a given set of pairs of atomic coordinates and subtomograms to learn their relationships in order to predict unknown atomic coordinates from a previously unseen set of subtomograms. The proposed regressor encoder–decoder architecture combines 3D convolutional extraction of features from subtomograms with a multilayer perceptron to predict Cartesian all-atom coordinates. Experiments on synthetic datasets show that DeepMDTOMO achieves low errors of coordinate prediction in presence of noise, MW, and large continuous conformational variability. Additionally, fine-tuning to a new motion demonstrates that learned representations capture general structure–density relationships rather than specific patterns. The results presented are encouraging and motivate future studies on speeding up subtomogram flexible-fitting methods with DL for fast atomic-scale conformational landscape determination from cryo-ET data.

## I. Introduction

Understanding structural dynamics of biomolecular complexes is essential for understanding their biological functions. Many complexes do not operate on a single rigid configuration but undergo continuous conformational changes that enable processes such as chromatin remodeling, gene regulation, and enzymatic activity. Capturing structural variability at the atomic scale remains a major challenge, particularly when studying macromolecules in their native cellular environment. [1]

Cryo-electron tomography (cryo-ET) has emerged as a powerful technique for *in situ* structural biology, enabling three-dimensional (3D) imaging of macromolecular complexes directly inside cells. In cryo-ET, individual biomolecular complexes (also called particles) are extracted from tomographic reconstructions (called tomograms) into individual volumes (called subtomograms) that naturally capture conformational heterogeneity. However, structural analysis of subtomograms is difficult due to a low signal-to-noise ratio (SNR) and missing-wedge (MW) artifacts. In classical cryo-ET workflows, the SNR is increased and the MW attenuated using iterative classification, alignment, and averaging of subtomograms [2] [3]. However, the classical workflows do not allow for capturing subtle conformational differences of the particles, as they can be hidden in the class averages. Recently, deep learning (DL) methods have been proposed to recover conformations in individual subtomograms, without averaging, which also allows for extracting continuous conformational variability from the data (gradual conformational changes with multiple intermediate conformational states), but these methods are solely based on cryo-ET data and result in low-resolution density-map representation of the particles [4] [5].

To address this problem, physics-based approaches have been developed, which integrate molecular modeling into experimental data analysis. They interpret structural differences in subtomograms by explicitly modeling molecular flexibility. Among them, methods based on normal mode analysis (NMA) provide a compact and physically meaningful description of large-scale collective motions. In the context of cryo-ET, HEMNMA-3D [6] was introduced to estimate poses (three Euler angles and shifts in the x, y, and z directions) and conformations of particles directly from subtomograms, by flexible fitting of an available starting conformational model into each individual subtomogram. In the NMA-based approach, the conformational parameters are amplitudes of a linear combination of normal modes (vectors showing directions of harmonic motions of atoms of different frequencies).

To improve physical accuracy of estimated conformational models, this framework was further extended with MDTOMO [7], which performs flexible fitting of subtomogram using classical molecular dynamics (MD) simulations (based on Newton’s equations of motion). MDTOMO allows obtaining atomic models of higher quality than HEMNMA-3D, but at the price of high computational cost.

To speed up MDTOMO, we are developing a supervised DL approach (referred to as DeepMDTOMO) that uses an encoder-decoder architecture [8] and learns the relationships between a given dataset of subtomograms and their corresponding Cartesian atomic coordinates (obtained with MDTOMO by analyzing the subtomograms) in order to predict the atomic models from a large subset of subtomograms unseen during the training. The encoder consists of stacked convolutional blocks and extracts spatial features from the given subtomograms, and the decoder is a multilayer perceptron (MLP) that maps the subtomogram feature vectors to the given atomic models (x, y, z coordinates of atoms). The problem is formulated as a regression from 3D subtomogram data to atomic coordinates. This design is inspired by DeepHEMNMA [9]. However, DeepHEMNMA learns to predict a small number of normal-mode amplitudes (usually, 5-10) per particle, whereas DeepMDTOMO aims to learn a large number of atomic coordinates (three times the number of atoms, with typical proteins consisting of tens of thousands of atoms). Also, DeepHEMNMA operates on 2D images, whereas DeepMDTOMO operates on MW-affected volumetric data.

In this article, we describe the concept of DeepMDTOMO and show the performance of the DL part of the framework using synthetic data that provide controlled ground-truth for training and evaluation of inference. To decouple the errors of MDTOMO from those of the DL model, we evaluate the DL performance for the case where the training is performed using exact ground-truth atomic coordinates instead of using MDTOMO-derived atomic coordinates (estimated from the synthetic subtomograms).

## II. Methods

The concept of DeepMDTOMO, based on MDTOMO and a DL neural network, is shown in Figure 1. In this section, we briefly recall MDTOMO and describe the DL network implementation.

**Figure 1.**
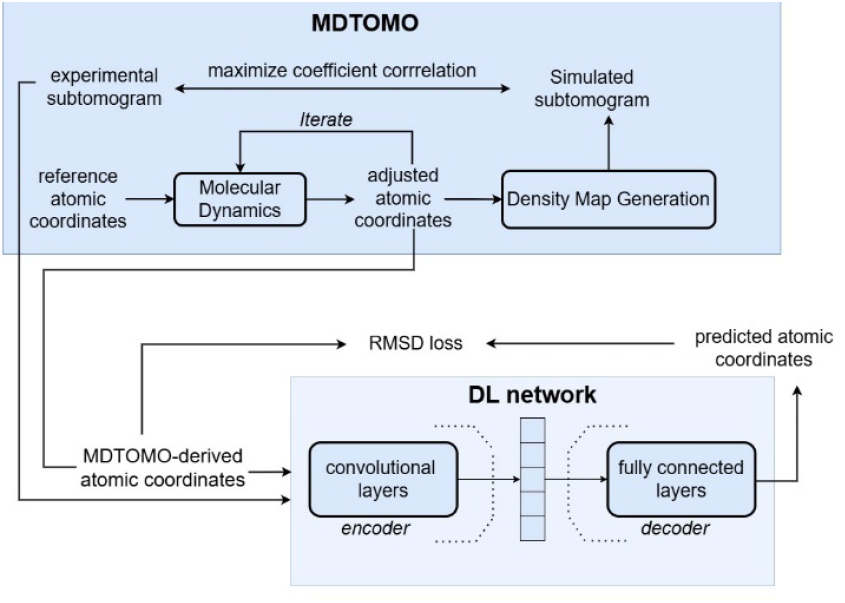
Concept of DeepMDTOMO. The inputs to MDTOMO are a reference atomic structure and subtomograms. The atomic structure is iteratively modified based on MD simulations until the corresponding simulated density map is aligned with the subtomogram. The MDTOMO-derived atomic coordinates and corresponding subtomograms are the inputs to the DL network that is trained by minimizing RMSD loss between predicted and MDTOMO-derived coordinates.

### A. MDTOMO

MDTOMO [7] is a computational framework designed to extract atomic-level biomolecular conformations from cryo-ET subtomograms. MDTOMO displaces atoms of a reference structure to match individual subtomograms, and the displacement is performed using classical MD simulations. The reference structures are typically obtained from the Protein Data Bank (PDB) [10]. This procedure captures both global collective motions and local atomic rearrangements, enabling a detailed description of continuous conformational variability.

### B. DL neural network

The DL network consists of an encoder and a decoder. The encoder consists of three convolutional blocks with channel dimensions (32, 64, 128), which extract spatial features from the input subtomograms (containing information on the particle’s conformation and pose). The encoded representation is compressed into a latent vector of dimension 256, which contains a low-dimensional description of the subtomograms. The decoder is implemented as a 3-layer multilayer perceptron with hidden dimensions (512, 1024, 2048), followed by a linear output layer that predicts the Cartesian coordinates (x, y, z) of all atoms for the chosen complex.

The training is formulated as a supervised regression problem, where the network learns to predict the Cartesian atomic coordinates corresponding to a given subtomogram. The model parameters are optimized by minimizing the loss that is the root-mean-square deviation (RMSD) between the predicted and ground-truth atomic coordinates. This loss function directly penalizes deviations in atomic positions and provides an intuitive and physically meaningful measure of structural accuracy.

## III. Experiments

### A. Data

Our simulation setting differs from a practical cryo-ET setting. While real tilt images contain many molecules that need to be extracted into individual volumes (subtomograms) from a tomogram reconstructed from these tilt images, we simulate sub-tilt images (a single molecule per tilt-series), which allows for a more direct and efficient simulation of the subtomograms (by a direct 3D reconstruction from the simulated images). Importantly, at the level of single molecules, the MW produces the same effects in both settings, as the MW effects are linked to the orientation of the tilt axis, which is the same in both settings (the tilt axis is perpendicular to the electron beam).

Synthetic cryo-ET subtomograms were generated using the ContinuousFlex [11] plugin for Scipion [12] software package, starting from the atomic structure of the chain A of adenylate kinase (AK) from the Protein Data Bank, PDB:4AKE, which will be referred to as reference structure. The first step of data synthesis was NMA of the reference structure to compute normal modes (vectors showing directions of harmonic motions of different frequencies). Low-frequency normal modes describe large-scale, collective motions of atoms relevant to biomolecular functions. Three lowest-frequency non-rigid normal modes (modes 7-9, Figure 2a) were used to simulate motion of AK at the next step of data synthesis (the first six normal modes describe rigid-body motions and are not used for simulating conformational states).

**Figure 2.**
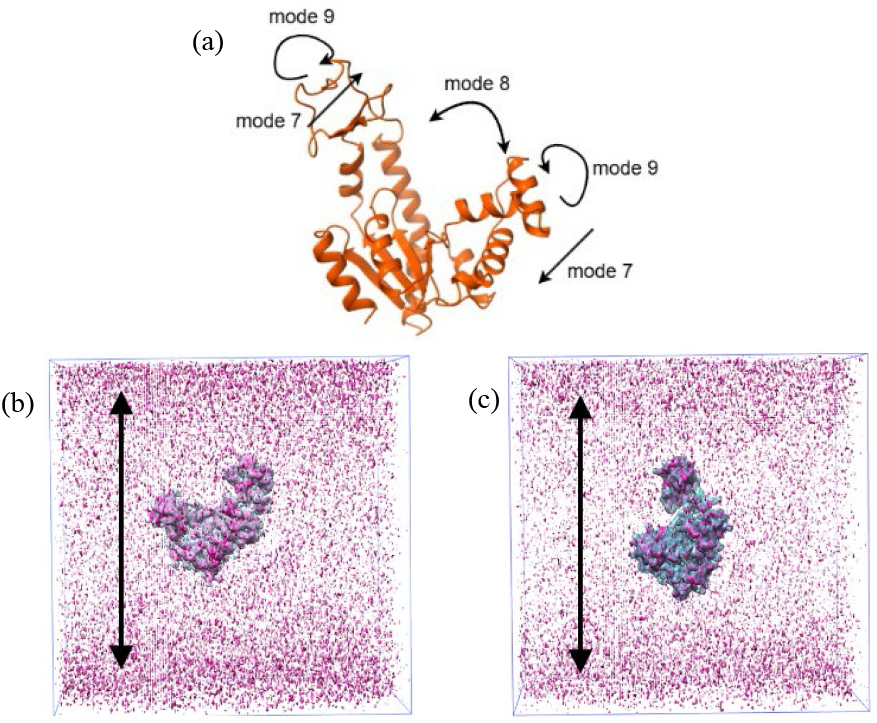
Atomic model of chain A of adenylate kinase (AK) and normal modes used for data synthesis, together with examples of synthetized subtomograms. (a) Atomic model of AK and motion directions for normal modes 7-9. (b,c) Overlap of a subtomogram (with noise, CTF, and MW) in pink and the corresponding ideal volume (no noise, CTF or MW) in cyan, for two examples of particles. The arrows indicate MW-induced elongations (particle and noise distortions).

Two sets of conformational states were generated: one using modes 7 and 8, the other using mode 9. A continuous set of elastically deformed atomic structures was obtained by sampling normal-mode amplitudes in the range [-120, 120]. The obtained atomic coordinates were used as the ground-truth for supervised training of the proposed DL network in the next subsection.

Each deformed atomic structure was randomly rotated and shifted (uniform distribution for the three Euler angles and three shifts) and c onverted into a 3D volume (density map) of size 128×128×128 voxels and a voxel size of (1.1Å)^3^. Here, these volumes will be referred to as ideal or noise-free volumes.

At the next step of data synthesis, each ideal volume was rotated (tilted) around the same rotation (tilt) axis (an axis perpendicular to the simulated electron beam, as in real settings) and, for each tilt angle, a 2D projection was calculated on a static image plane. The tilting was performed in the tilt-angle range [−60^°^,+60^°^] with an increment of 1^°^, as is often the case in cryo-ET. The projections were further degraded by CTF effects (defocus of 0.5 μm and spherical aberration of 2 mm) and noise (SNR of 0.1), mimicking realistic cryo-ET imaging conditions. 3D reconstructions (volumes of size 128×128×128 voxels) were then obtained from these images. Here, these volumes will be referred to as noisy volumes although they are not only degraded by noise but also by MW and CTF.

The MW can be observed in Fourier space as a wedge-shaped empty region related to the missing tilt images (tilt-angle range beyond [−60°, +60^°^]). The MW-induced deformations on the volumes in real space can be observed in Figure 2b,c that shows MW-induced elongations in the direction perpendicular to the tilt axis. Generally speaking, the analysis of MW-affected subtomograms presents difficulties due to a combination of the two types of molecular shape deformations: those due to continuous conformational changes and those due to MW.

In total, 40,000 subtomograms were generated for conformations based on a linear combination of normal modes 7 and 8. Among those, one half (20,000 subtomograms) is noise-free (ideal), and each noisy subtomogram has a corresponding ideal-volume counterpart representing the exact same atomic coordinates. Additionally, 40,000 subtomograms were generated for conformations based on normal mode 9, among which, again, 20,000 noise-free (ideal) and 20,000 noisy, where each noisy subtomogram has an ideal counterpart corresponding to the same underlying conformational state, orientation, and position of AK. Each synthetic subtomogram is paired with its corresponding ground-truth atomic coordinates, enabling supervised training and quantitative evaluation of inference with the proposed DL network. As shown in the next subsection, the subtomograms generated using normal modes 7 and 8 were used to evaluate the performance of training and inference with the DL network, whereas those generated using normal mode 9 served to evaluate the potential of the network to generalize and fine-tune on different conformational distributions than the one used for the initial training.

### B. Progressive training procedure, validation, and inference

The model training was conducted in two consecutive steps to progressively improve performance and robustness. In step 1, the network was trained using only noise-free data generated based on normal modes 7 and 8 (learning rate 0.01, a cosine decay schedule). The use of noise-free data for training in this step allowed to learn the mapping between subtomograms and ground-truth atomic coordinates under ideal conditions. The inference in this step was performed using noise-free subtomograms generated using normal modes 7 and 8 (ground-truth atomic coordinates were only used for evaluation of the inference results). The dataset for this step (20,000 noise-free subtomograms and corresponding ground-truth atomic coordinates) was split as follows: 16,000 for training, 2,000 for validation, and 2,000 for inference.

In step 2, again using data based on normal modes 7 and 8, training was performed using a balanced, 50%-50% mix of 8,000 noise-free and 8,000 noisy subtomograms, while the validation set consisted exclusively of noisy subtomograms (2,000 noisy subtomograms). In this step, the model was initialized with the learned weights from step 1 (learning rate 0.01, a cosine decay schedule), allowing it to retain knowledge of clean conformational mappings while adapting to realistic cryo-ET noise. The inference dataset in step 2 consisted of 2,000 noisy subtomograms that were different from those used for validation.

Finally, we used transfer learning to evaluate the model’s ability to generalize to previously unseen conformations, namely those generated based on normal mode 9, which was not included in the original training data (steps 1-2). Subtomograms based on normal mode 9 and the corresponding ground-truth atomic conformations were used to fine-tune the model initialized with the learned weights from step 2 (learning rate 0.01, a cosine decay schedule). The model was fine-tuned using 8,000 noisy and 8,000 noise-free subtomograms for training. The validation and inference datasets contained only noisy data, as in step 2 (2,000 noisy subtomograms for validation and a different set of 2,000 noisy subtomograms for inference).

### C. Results

The results presented in Figure 3 and Table 1 indicate that such a progressive training strategy (referred to as case 1) allows the network to progressively learn from clean data, adapt to noisy conditions, and extend its predictive capability to unseen features (conformational variability based on mode 9) while leveraging previously learned features (from conformational variability based on normal modes 7 and 8).

**Table 1:** Inference performance under different training strategies, noise percentages, and normal-mode settings. Results are reported on the inference data sets in terms of RMSD statistics between predicted and ground-truth atomic coordinates (mean ± standard deviation, minimum, maximum, median) and number of predictions with RMSD < 3 Å. Case 1 shows the progressive training and transfer to an unseen conformational mode (mode 9), while Case 2 reports performance for an independently trained model on modes 7 & 8 only on noisy data.

|  | Percentage of noisy subtomograms |  |  | Normal Modes | RMSD statistics at inference |  |  |  |  |
| --- | --- | --- | --- | --- | --- | --- | --- | --- | --- |
| | Train | Validation | Inference | | Mean ± std (Å) | Min (Å) | Max (Å) | Median (Å) | <3Å |
| Case 1: Step 2 | 50% | 100% | 100% | 7 & 8 | 1.63 ± 0.48 | 0.63 | 4.37 | 1.56 | 1962 |
| Case 1: Transfer | 50% | 100% | 100% | 9 | 1.63 ± 0.48 | 0.68 | 4.39 | 1.58 | 1974 |
| Case 2 | 100% | 100% | 100% | 7 & 8 | 3.38 ± 1.00 | 1.24 | 8.31 | 3.24 | 808 |

**Figure 3.**
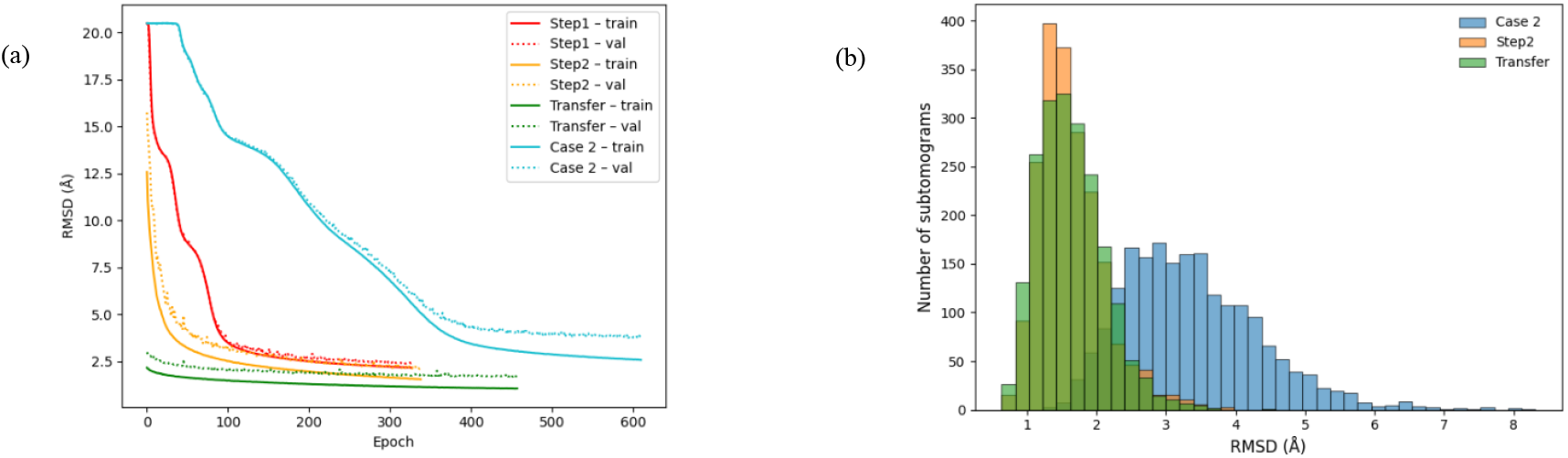
Training and inference performance. (a) Training and validation RMSD loss curves across the two training steps, showing progressive improvement from noise-free training to noise aware training and transfer learning on a new conformational mode, as well as training from noisy data alone (referred to as case 2). (b) Distribution of RMSD values between predicted and ground-truth atomic coordinates at inference step, highlighting a shift in accuracy across training.

Training on noise-free (ideal) subtomograms (step 1) provides a stable initialization and allows the network to capture the geometric relationship between volumetric densities and atomic coordinates without the confounding effect of noise, MW, and CTF (Figure 3a).

Introducing noisy subtomograms in step 2 induces stable training, which is indicated by a stable convergence behavior observed in the loss curves in Figure 3a. For comparison, we also show the results obtained for training on noisy data alone, in a single step (referred to as case 2). In case 2, the dataset of 20,000 noisy subtomograms was split into 16,000 for training, 2,000 for validation, and 2,000 for inference. Figure 3 and Table 1 show that mixing clean and noisy data during training (step 2) performs better than noisy-only training (case 2), yielding a lower mean RMSD (1.63 Å vs. 3.38 Å) and a higher number of accurate predictions (1,962 vs. 808 out of 2,000 with RMSD < 3 Å).

Transfer learning succeeded, enabling the model to generalize to previously unseen conformational variability, and produced a low mean RMSD (1.63 Å) and predictions with RMSD < 3 Å for 1,974 (out of 2,000) inference samples (Figure 3, Table 1).

The presented results demonstrate, under controlled synthetic conditions, that the proposed DL network for DeepMDTOMO can learn a direct regression from cryo-ET subtomograms to atomic coordinates. The progressive training strategy plays a central role in the observed performance gains, enabling the model to first learn the idealized mapping under noise-free conditions and then gradually adapt to realistic noisy data and previously unseen conformational variability.

An important aspect of the transfer-learning results presented here is their relevance for future application to experimental cryo-ET data. In experimental conditions, the underlying conformational coordinates and dominant motion modes are unknown, and it is tempting to use simulation-based training and then fine-tune the trained model on experimental data, but one cannot assume that observed particles follow the simulated conformational variability (here, using normal modes) that was used for simulation-based training. Fine-tuning and evaluating the model on data generated using a normal mode unseen in initial training therefore provides a practical test of the model’s capacity to generalize beyond the specific deformation subspace used for initial training. The strong performance obtained after adaptation to mode 9 suggests that the learned representation captures general structure–density relationships rather than mode-specific patterns, which is encouraging for future deployment on experimental datasets.

### D. Speed

Using 16,000 subtomograms for training and 2,000 for validation took approximately 48 hours of wall-clock time on a single NVIDIA V100 GPU (16 GB). Inference on 2,000 subtomograms required 2.5 minutes on a single NVIDIA RTX 4500 Ada GPU.

## IV. Conclusion

In this work, we proposed a DL neural network architecture for DeepMDTOMO, a supervised deep learning framework for predicting atomic coordinates of biomolecular complexes directly from cryo-ET subtomograms. The method learns a regression from volumetric data to Cartesian atomic positions. By combining a convolutional encoder with a multi-layer perceptron decoder, the proposed approach is able to handle the high-dimensional output space associated with high conformational variability of atomic structures and a large number of atoms.

Experiments on realistic synthetic cryo-ET data demonstrate that the DL network achieves accurate structure prediction, with a mean RMSD error of 1.63Å relative to ground-truth atomic models. A progressive training strategy, starting from noise-free data, extending to mixed noisy conditions, and followed by transfer learning, was shown to significantly improve training robustness and inference generalization. In particular, successful adaptation to conformations generated from an unseen normal mode indicates that the initially learned representation is not restricted to a single deformation subspace. This property is especially important for potential applications of simulation-based training to experimental cryo-ET data, where the underlying conformational modes are unknown.

Overall, the results support the feasibility of using supervised DL as a fast approach to speed up computationally intensive physics-based flexible fitting methods, such as MDTOMO, which will be demonstrated in the future. Specifically, this article shows the performance of a DL network for DeepMDTOMO framework using simulated ground-truth atomic coordinates for supervised training, while the future work will address training using atomic coordinates estimated with MDTOMO from subtomograms.

The proposed framework opens the way toward scalable atomic-level conformational analysis from cryo-ET data. Future work will focus on scaling to larger macromolecular complexes (*e*.*g*., nucleosome, ribosome, etc.) and validation with experimental datasets.

## Acknowledgment

We acknowledge the support of the ANR (ANR-23-CE45-0012-03) and access to HPC resources of CINES and IDRIS granted by GENCI (AD010710998R3, AD010714089R1).

## Notes

### Competing Interest Statement

The authors have declared no competing interest.

